# Lack of evidence for anthocyanins contributing to pigmentation of *Chenopodium quinoa*

**DOI:** 10.64898/2026.04.07.717023

**Authors:** Luna Theresa Lingemann, David Biley, Jakob Maximilian Horz, Najnin Khatun, Boas Pucker

## Abstract

While most plant lineages are pigmented by anthocyanins, several families in the Caryophyllales represent a major exception by showing a replacement of anthocyanin pigmentation by betalain pigmentation. The mutual exclusion of anthocyanins and betalains at the family level has been well established for over 50 years and has been mechanistically explained. Chenopodiaceae are a betalain-pigmented lineage lacking a key anthocyanin biosynthesis gene and lacking the key activating transcription factor of the anthocyanin biosynthesis.

A publication by Zhang et al., 2024 claims that anthocyanins would be responsible for the red pigmentation in leaves of *Chenopodium quinoa*. Here, we assessed this study and reanalyzed the RNA-seq datasets generated in this study to demonstrate that there is no evidence for anthocyanin biosynthesis, but activity of the betalain and carotenoid biosynthesis could explain the observed pigmentation of quinoa leaves.

## Introduction

Pigmentation of plants has fascinated plant scientists for almost 200 years (Marquart, 1835; Mendel, 1865). Responsible for the colors of most plant lineages are anthocyanins, a group of specialized metabolites with diverse chemical modifications conferring orange, red, pink, purple, blue, or almost black hues (Winkel-Shirley, 2001; Grotewold, 2006; Grünig *et al*., 2025). Anthocyanins fulfill a range of ecological functions including pollinator and seed disperser attraction, protection of leaves as sunscreen, camouflage, and mimicry (Grünig *et al*., 2025). Numerous ecological and physiological functions might explain why anthocyanins are spread across almost all land plant lineages. Major exceptions are the Cucurbitaceae (Choudhary *et al*., 2026) and several families in the Caryophyllales (Timoneda *et al*., 2019). Many families within the Caryophyllales are able to produce a different type of pigment, betalains, which appear mutually exclusive with anthocyanin pigmentation (Kimler *et al*., 1970; Tanaka *et al*., 2008; Brockington *et al*., 2011; Timoneda *et al*., 2019). Plant families in the Caryophyllales are pigmented by anthocyanins or betalains, but the two pigments have not been observed together in the same plant in nature (Tanaka *et al*., 2008; Timoneda *et al*., 2019). However, heterologous expression of anthocyanin biosynthesis genes in betalain-pigmented species or vice versa has demonstrated that it is possible to produce both pigments in the same plants (He *et al*., 2020; Sakuta *et al*., 2021). The complex pigment evolution in the Caryophyllales resulting in the mutual exclusion of anthocyanins and betalains has been explained by at least four independent events of betalain biosynthesis emergence (Sheehan *et al*., 2020) and corresponding losses of the anthocyanin biosynthesis (Pucker *et al*., 2024). A minimal betalain biosynthesis typically involves three enzymes including tyrosine hydroxylase (CYP76AD1), L-DOPA 4,5-dioxygenase (DODAa), and a glucosyltransferase (Vogt, 2002; Christinet *et al*., 2004; Tanaka *et al*., 2008; Brockington *et al*., 2015; Timoneda *et al*., 2019). Anthocyanins are formed by one branch of the flavonoid biosynthesis involving the somewhat committed enzymes dihydroflavonol 4-reductase (DFR), anthocyanidin synthase (ANS), anthocyanin-related glutathione S-transferase (arGST), and UDP-dependent anthocyanidin 3-O-glucosyltransferase (UGT) (Grünig *et al*., 2025). Since the required genes of both pigment biosynthesis pathways are known, it is possible to check on the genetic level whether a plant would be able to produce one of these pigments. If a crucial gene is missing or not expressed, a biosynthesis route would be blocked. This has been demonstrated numerous times for the anthocyanin biosynthesis across many plant lineages (Ho & Smith, 2016; Wheeler *et al*., 2022; Marin-Recinos & Pucker, 2024).

There have been studies claiming the detection of anthocyanins in pitaya, a plant species belonging to a clearly betalain-pigmented lineage, but these claims have been refuted by exploring the presence and activity of biosynthesis genes (Pucker *et al*., 2021; Khan, 2022; Pucker & Brockington, 2022). To the best of our knowledge, there is currently no solid evidence for the claim that anthocyanins and betalains would contribute to pigmentation in the same plant species (Pucker & Khan, 2026).

Here, we are addressing a recent study by Zhang et al., 2024, which claims that anthocyanins would be responsible for red pigmentation of *Chenopodium quinoa* leaves (Zhang *et al*., 2024). We provide evidence that the anthocyanin biosynthesis is blocked by lack of a crucial gene and lack of detectable expression of another gene. As alternative explanations for the observed red color in quinoa leaves, we propose betalains and carotenoids by showing the expression patterns of corresponding biosynthesis genes.

## Results & Discussion

The study by Zhang et al., 2024 claims that anthocyanins would be responsible for the red pigmentation of *C. quinoa* leaves, but fails to provide solid evidence and lacks details regarding the applied methods and analyzed datasets.

### Genetic factors suggest blocks in the anthocyanin biosynthesis

The biosynthesis of anthocyanins in *C. quinoa* is probably fully blocked by the lack of arGST (**Fig. 1**). While it is impossible to prove the absence of a gene, a microsynteny analysis indicates the absence of an arGST in the Amaranthaceae/Chenopodiaceae and genome-wide searches revealed no evidence of arGST. These results align with a previous analysis reporting an arGST loss as a potential explanation for the evolutionary transition from anthocyanin to betalain-pigmentation in multiple Caryophyllales lineages (Pucker *et al*., 2024). Our analysis was restricted to orthologs of *A. thaliana TRANSPARENT TESTA 19* (*TT19*) and *P. hybrida Anthocyanin 9* (*An9*), because orthologs of *BRONZE2* (*Bz2*) have only been discovered in Poaceae (Khatun *et al*., 2025).

**Fig. 1:**
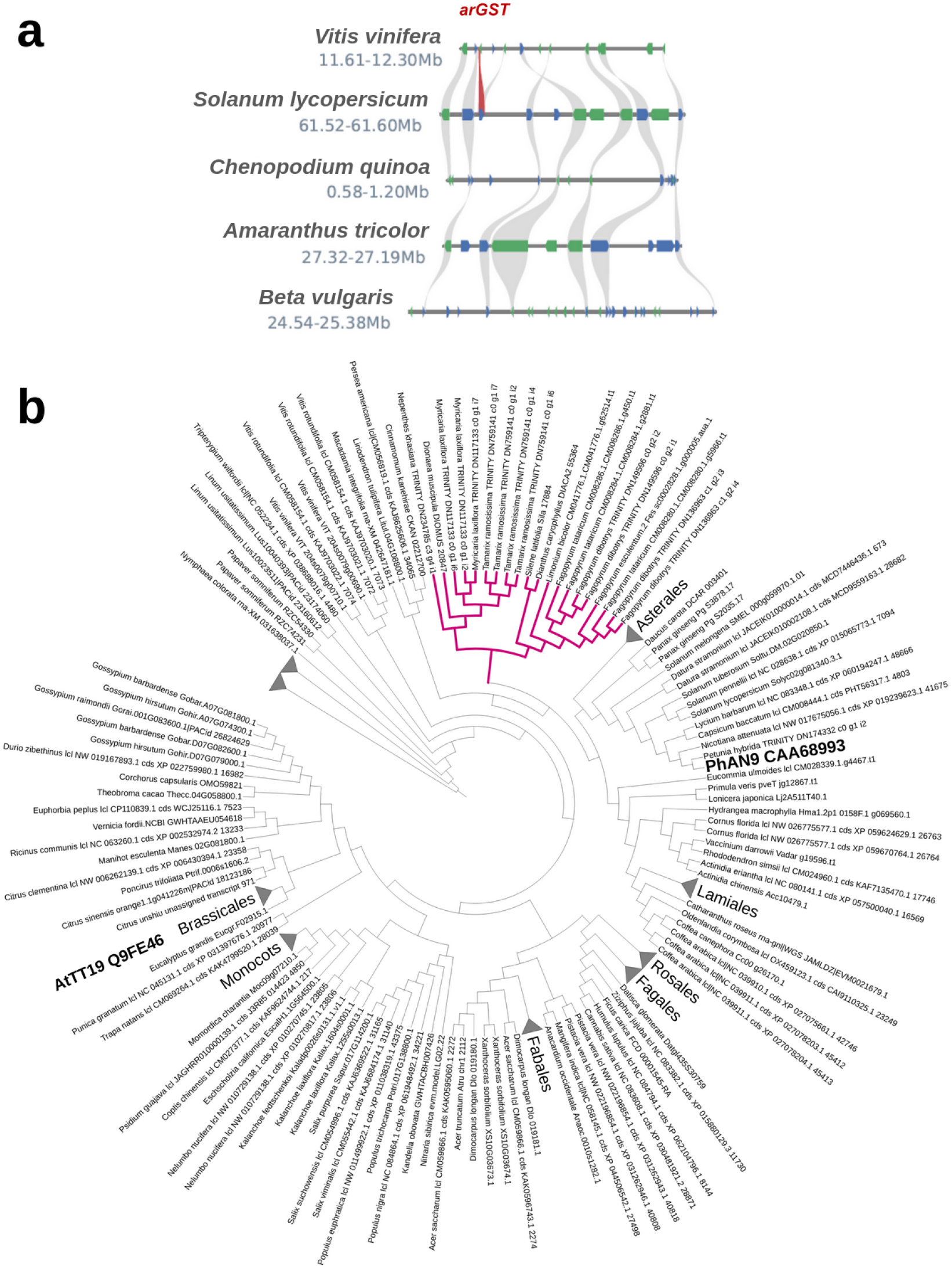
Microsynteny analysis around the arGST locus in anthocyanin-pigmented species suggests absence of a corresponding arGST gene in Amaranthaceae/Chenopodiaceae (a). A phylogenetic analysis of arGST-like sequences does not reveal any candidate in a betalain-pigmented Caryophyllales species (b). Only anthocyanin-pigmented Caryophyllales species (magenta) show orthologs of arGST. The gene tree with expanded arGST clades is available as Additional file B.

The anthocyanin biosynthesis is usually controlled by a complex of three transcription factors: a MYB, a bHLH, and a WD40 protein (Gonzalez *et al*., 2008; Lloyd *et al*., 2017). The MYB component of this complex has been frequently observed to be the most specific factor in controlling the anthocyanin biosynthesis (Marin-Recinos & Pucker, 2024). However, the orthologous MYB in sugar beet, a close relative of quinoa, has been reported to have lost the ability to form this MBW complex for anthocyanin biosynthesis activation (Pucker *et al*., 2024) and was instead co-opted by the betalain biosynthesis (Hatlestad *et al*., 2015). An inspection of the MYB candidate sequences in quinoa revealed no clear ortholog (Additional file H). It remains unclear whether this is an annotation artifact in the particular dataset or a general feature of quinoa. Given the apparent absence of an orthologous MYB in quinoa there is probably no formation of a MBW complex and consequently no activation of the anthocyanin biosynthesis genes.

### Gene expression patterns show evidence for betalain and carotenoid biosynthesis, not anthocyanin biosynthesis

Our reanalysis of the anthocyanin biosynthesis gene expression in *C. quinoa* based on the RNA-seq datasets published by Zhang et al., 2024 revealed no evidence for transcription of *DFR* and very low transcription of *ANS* (**Fig. 2a**). The residual activity observed for *ANS* could be explained by an ongoing proanthocyanidin biosynthesis as this branch of the flavonoid biosynthesis appears to be active in quinoa - at least in dark pigmented grains (Wang *et al*., 2020). The relatively high activity of *FLS*, which encodes an enzyme competing with the anthocyanin biosynthesis, supports the notion that no relevant anthocyanin biosynthesis can take place. The betalain biosynthesis genes appear to be slightly more active, but their activity does not explain the reported pigmentation differences (**Fig. 2b**). A previous study reported betalains as the major contributor to red quinoa coloration, but noticed a discrepancy between expression of known betalain biosynthesis genes and pigment accumulation which was attributed to other factors (Feng *et al*., 2023). While the carotenoid biosynthesis appears to be slightly more active, there is also no clear difference between samples that were labeled as green and those that were labeled as red (**Fig. 2c**). In summary, none of the pigment biosynthesis pathways that could result in reddish pigmentation appears to explain the reported color difference. This apparent contradiction of our analysis results and the results reported by Zhang et al., 2024 might be explained by a mismatch between the sample description presented in their manuscript (day 70, 90, 110) and the samples provided via the Sequence Read Archive (day 5, 20, 35).

**Fig. 2:**
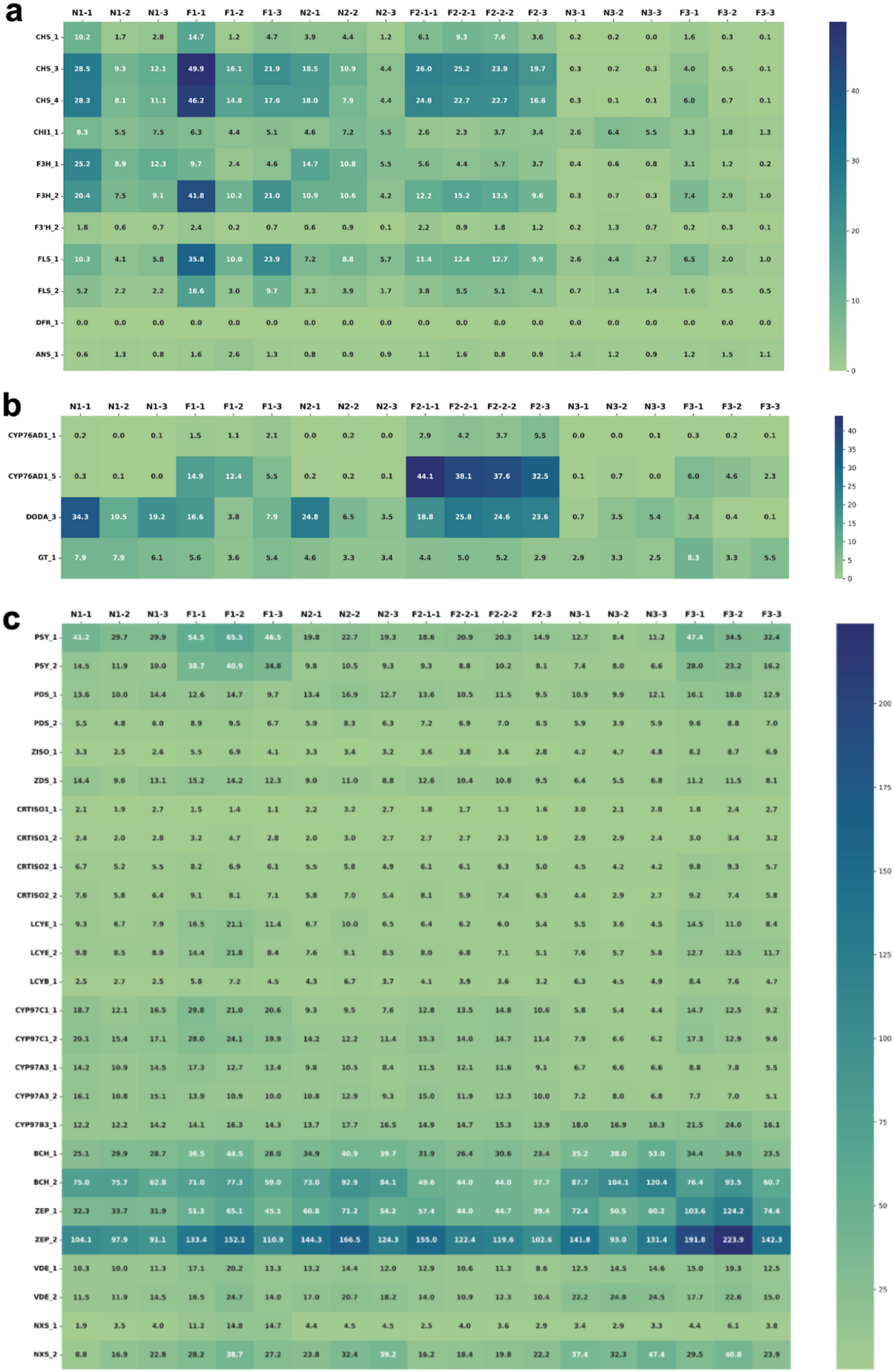
Expression analysis for genes involved in the anthocyanin (a), betalain (b), and carotenoid (c) biosynthesis. Chalcone synthase (CHS), chalcone isomerase (CHI), flavanone 3-hydroxylase (F3H), flavonoid 3’-hydroxylase (F3’H), flavonol synthase (FLS), dihydroflavonol 4-reductase (DFR), and anthocyanidin synthase (ANS) are enzymes associated with the anthocyanin biosynthesis. CYP76AD1, L-DOPA 4,5-dioxygenase (DODA), UDP–glucose: cyclo-DOPA 5-O-glucosyltransferase (GT) are enzymes in the betalain biosynthesis. Phytoene synthase (PSY), phytoene desaturase (PDS), ζ-carotene isomerase (Z-ISO), ζ-carotene desaturase (ZDS), carotenoid isomerase (CRTISO), lycopene ε-cyclase (LCYE), lycopene β-cyclase (LCYB), cytochrome P450-type monooxygenases 97A, 97B and 97C (CYP97A3/B3/C1), β-carotene hydroxylase (BCH), violaxanthin de-epoxidase (VDE), zeaxanthin epoxidase (ZEP), and neoxanthin synthase (NXS) form the carotenoid biosynthesis. Candidate genes with a combined expression <5 TPM were excluded from the main figure, but are displayed in Additional file E.

Re-analysis displayed significant enrichment in phenylpropanoid- and flavonoid-biosynthesis pathway expression (**Fig. 3a**). A look at the annotation of specific transcripts for enzymes in the flavonoid-pathway (**Fig. 3b**) reveals the differential expression is limited to *TT4* (chalcone synthase), *TT5* (chalcone/flavanone isomerase), *FLS1, F3H* and *ANS*, for the latter of which the log2 fold change is comparatively low. *DFR* as the key enzyme for the transition into the anthocyanin-biosynthesis pathway is notably absent. Other enriched pathways include pyrimidine metabolism, tropane, piperidine and pyridine alkaloid biosynthesis, cyanoamino acid metabolism, sesquiterpenoid and triterpenoid biosynthesis of which none are mentioned in the original publication by Zhang et al. 2024.

**Fig. 3:**
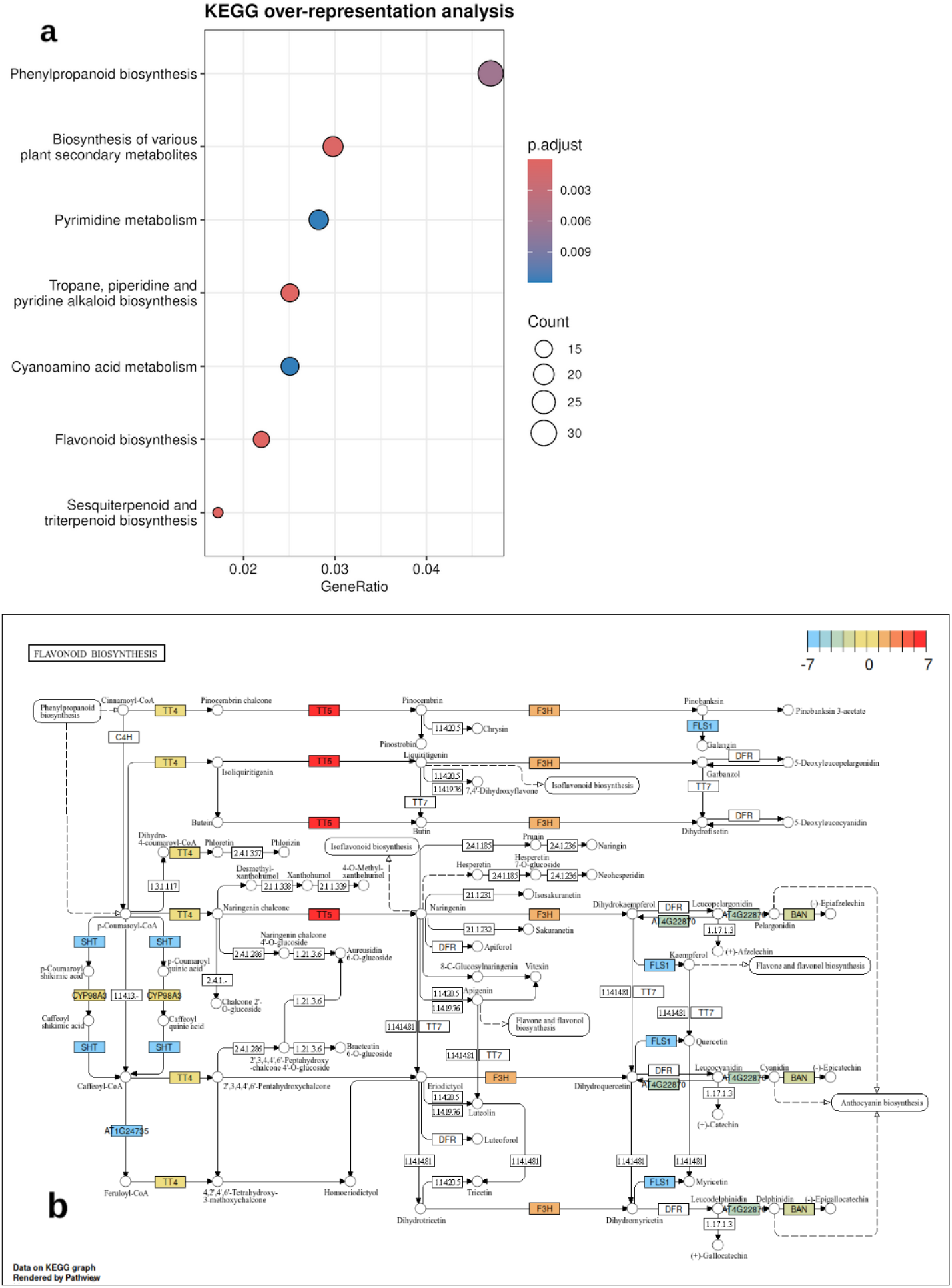
Dotplot of the over-representation analysis of KEGG-pathways (Kanehisa & Goto, 2000) with differentially expressed genes between the two sampled varieties (M202 and M146). The adjusted p-value of the pathway’s enrichment is displayed by the color gradient, the number of enriched genes by the size of the dots and the enrichment factor by the position on the x-Axis (a). Differential expression of enzymes mapped onto the KEGG anthocyanin pathway based on annotations from *Arabidopsis thaliana* (ath00941). The enzymes are colored on a gradient from red at log2FC cutoff = 7 to blue at log2FC cutoff = -7 where |log2FC| close to 1 appear yellow, while enzymes with |log2FC| < 1 and enzymes with no expression are not colored at all (b).

### No strong metabolomics evidence for anthocyanins

It is important to emphasize that previous studies have provided solid evidence for the presence of betalains in quinoa (Escribano *et al*., 2017; Henarejos-Escudero *et al*., 2018; Steimker *et al*., 2026). In addition, screens for anthocyanins across multiple quinoa varieties have not been successful (Ong *et al*., 2020). While it is theoretically impossible to prove the absence of a metabolite group, these studies strongly suggest that anthocyanins are not relevant for the pigmentation in quinoa.

Zhang et al., 2024 determined the total anthocyanin content in leaf samples following a standard spectrophotometric method, by measuring the absorption of the extract at 530 nm and 657 nm (Serrano *et al*., 2012). However, this is not sufficient to prove anthocyanin identity, because other pigments like betalains also show absorption at similar wavelengths (Pucker & Khan, 2026). Many betalains have absorbance maxima between 532 and 550 nm (Khan & Giridhar, 2015). The absorbance values reflect the absorbances of the whole extract and not that of anthocyanins contained in the sample as was stated in the methods section of the paper by Zhang et al., 2024. The pigment content measured can most likely be attributed to betalains instead of anthocyanins due to aforementioned reasons.

The LC-MS/MS analysis presented by Zhang et al., 2024 has some limitations. Firstly, the methods section omits information on multiple important parameters and devices used in both the chromatography and mass spectrometry, including LC parameters such as column type, mobile phases, gradients or flow rate and injection volume. Key MS/MS acquisition parameters like ionisation mode (ESI+ or ESI-), precursor ions selected for fragmentation, and collision energies are also not reported. These details are essential to enable validation of the results by independent replication. Furthermore, neither the raw data, retention times or MS/MS spectra are shared in the results.

### Conclusion and recommendations for future studies

The mutual exclusion of anthocyanin and betalain pigmentation in the Caryophyllales is well supported by numerous studies on different levels thus future studies contradicting it would require very strong evidence (Pucker & Khan, 2026). Superficial transcriptomic or metabolomic analyses, sometimes conducted by external service providers, are a likely cause for issues. The widespread knowledge about anthocyanins and the dominance of anthocyanin biosynthesis genes in databases like Kyoto Encyclopedia of Genes and Genomes (KEGG) (Kanehisa *et al*., 2023) and Gene Ontology (GO)(Gene Ontology Consortium, 2021) might be explanations why enrichment analyses could generate the impression that anthocyanin biosynthesis would be active in betalain-pigmented lineages. In-depth investigations of well-annotated individual genes would help to reach well-supported conclusions.

## Methods

### Microsynteny analysis and phylogenetic analysis indicating absence of arGST

The genome assemblies and their corresponding annotation files for the five species, *Vitis vinifera* (Shi *et al*., 2023), *Solanum lycopersicum* (Shirasawa & Ariizumi, 2024), *Chenopodium quinoa* (Jarvis *et al*., 2017), *Amaranthus tricolor* (Wang *et al*., 2023), and *Beta vulgaris* (Sielemann *et al*., 2023), were retrieved from NCBI and Phytozome. Completeness of the assemblies was assessed with BUSCO v6.0.0 (Manni *et al*., 2021) [in protein mode] using the *embryophyta_odb12* lineage datasets. Microsynteny analysis was performed to visualize the conserved genomic regions using JCVI/MCscan (Tang *et al*., 2024). The 10 protein-coding genes flanking the *arGST* loci were selected to demonstrate the preservation of gene order. To ensure that a translocated arGST at a different genomic position would be discovered, we conducted a comprehensive analysis by searching with the *A. thaliana* (Q9FE46) (Kitamura *et al*., 2004) and *Petunia hybrida* (CAA68993) (Alfenito *et al*., 1998) arGST sequences using a customized Python wrapper script for BLASTp-based searches and integration of results (Pucker & Iorizzo, 2023). All collected arGST candidate sequences were aligned with MAFFT v7.505 (Katoh & Standley, 2013) using default parameters. Alignment columns with low occupancy were trimmed with the Python script algntrm using default parameters (Pucker & Iorizzo, 2023). A gene tree was inferred with IQ-TREE v3.1.0 (Wong *et al*., 2025) using the model finder for the identification of optimal parameters (Q.PLANT+R10 was identified as best fit model). The resulting tree file was visualized in iTOL (Letunic & Bork, 2024).

### Phylogenetic analysis of the anthocyanin, betalain, and carotenoid biosynthesis genes

All analyses were conducted based on the Cquinoa_392_v1.0 annotation of *C. quinoa* retrieved from Phytozome (Jarvis *et al*., 2017). Flavonoid biosynthesis and carotenoid biosynthesis genes were identified with KIPEs v3.2.6 (Rempel *et al*., 2023) based on the flavonoid biosynthesis dataset v3.4 and carotenoid biosynthesis dataset v2.0. MYB_annotator v1.0.3 (Pucker, 2022) was used to identify orthologous MYBs in Quinoa. The identification of candidates for DODA, CYB76AD1, and the betalain regulating MYB was conducted based on orthology to previously reported sequences. Briefly, sequences of DODA1α and CYP76AD1α (Brockington *et al*., 2015) and sequences of the betalain biosynthesis activating MYB (Stracke *et al*., 2014) served as bait in collecting initial candidates based on local sequence similarity assessed by BLASTp v2.12.0 as implemented in collect_best_BLAST_hits.py (Altschul *et al*., 1990; Pucker & Iorizzo, 2023). Global alignments were generated with MAFFT v7.505 (Katoh & Standley, 2013) and trimmed with the Python script algntrm using default parameters (Pucker & Iorizzo, 2023). IQ-TREE v3.1.0 (Wong *et al*., 2025) was used to infer gene trees by running the model finder followed by an analysis based on the best model and conducting 1000 rounds of bootstrapping. iTOL (Letunic & Bork, 2024) was used for manual inspection of the tree and the characterized sugar beet orthologs Bv_ptjc (DODA1α), Bv_ucyh (CYP76AD1α), and Bv_jkkr (betalain MYB) indicated the orthologs in quinoa (Additional file F, G, H).

### Gene expression analysis of anthocyanin, betalain, and carotenoid biosynthesis

The gene expression data linked from the study by Zhang et al., 2024 were retrieved from the Sequence Read Archive with fastq-dump (NCBI, 2020). It is important to note that the metadata at the SRA do not match the information provided in the manuscript. FASTQ files were processed with kallisto v0.44 (Bray *et al*., 2016) with the coding sequences of Cquinoa_392_v1.0 (Jarvis *et al*., 2017) serving as reference. Downstream processing with Python scripts was conducted as previously described (Pucker *et al*., 2024). Heatmaps for the visualization of gene expression patterns were generated with a customized Python script utilizing the packages seaborn (Waskom, 2021) and matplotlib (Hunter, 2007) and available via GitHub (https://github.com/PuckerLab/QuinoaPigmentation).

### Enrichment analyses

The peptide sequences of *C. quinoa* (GCF_001683475.1) (Jarvis *et al*., 2017) were used to find orthologs in the peptide sequences of the *Arabidopsis thaliana* TAIR10 assembly using Orthofinder 3.0.1b1 (Emms & Kelly, 2019) (Additional file D). PyDESeq2 v0.5.4 (Muzellec *et al*., 2023) was used for the differential expression analysis (Additional file C). R v4.4.2 was used with clusterProfiler (Yu *et al*., 2012) for over-representation analysis and pathview (Luo & Brouwer, 2013) to display the results of differential expression in KEGG-pathways. The environment was set up using the bioconductor docker image “bioconductor/bioconductor_docker:RELEASE_3_20”. The scripts for differential expression and enrichment analyses are available via GitHub.

## Declarations

### Ethics approval and consent to participate

Not applicable

## Consent for publication

Not applicable

## Availability of data and materials

Sequencing data are available via SRA/ENA (PRJNA986269) and the derived count table containing TPMs is included as Additional file A. Scripts developed for this study are available via https://github.com/PuckerLab/QuinoaPigmentation.

## Authors’ contributions

BP designed the research project and supervised the work. LL, DB, JH, NK, and BP conducted the analyses. LL, DB, JH, BP interpreted the results and wrote the manuscript. All authors approved the final version of the manuscript and agreed to its submission.

## Acknowledgements

This work was supported by the de.NBI Cloud within the German Network for Bioinformatics Infrastructure (de.NBI) and ELIXIR-DE (Forschungszentrum Jülich and W-de.NBI-001, W-de.NBI-004, W-de.NBI-008, W-de.NBI-010, W-de.NBI-013, W-de.NBI-014, W-de.NBI-016, W-de.NBI-022). We thank all members of the Plant Biotechnology and Bioinformatics group for their support and feedback during the process.

