## Supplementary figures and images for "Lack of evidence for anthocyanins contributing to pigmentation of *Chenopodium quinoa*"

### Additional file B

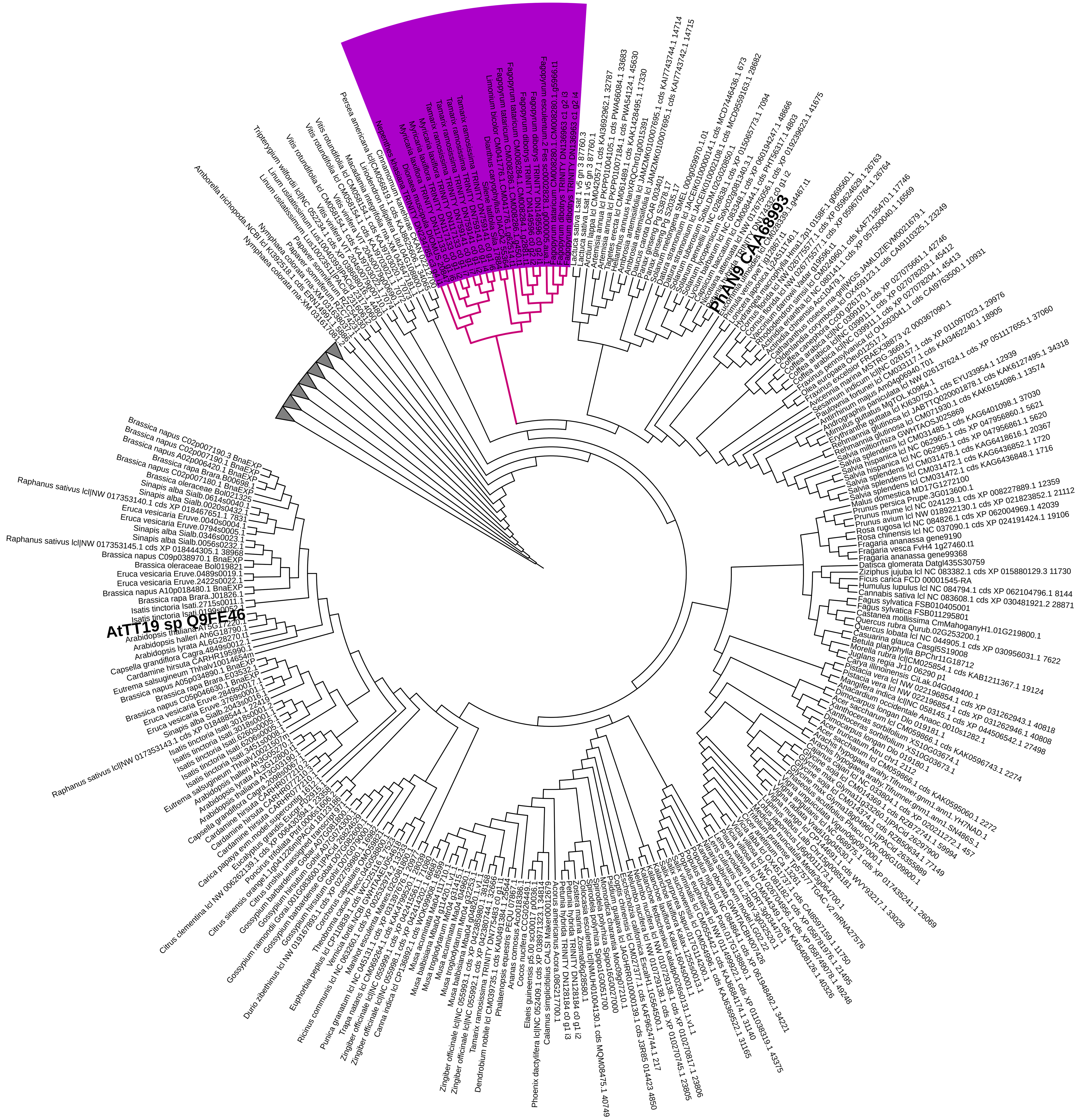

### Additional file F

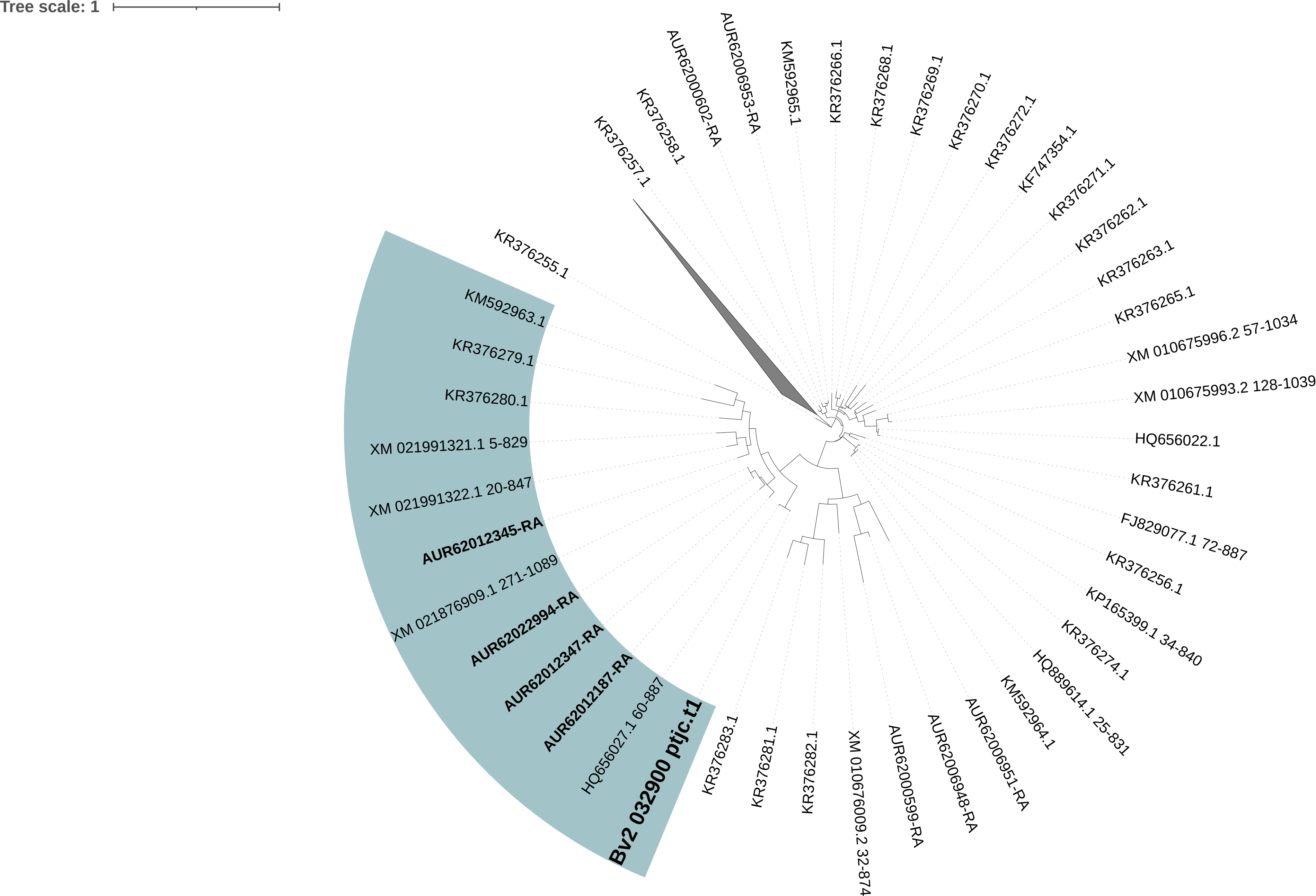

### Additional file G

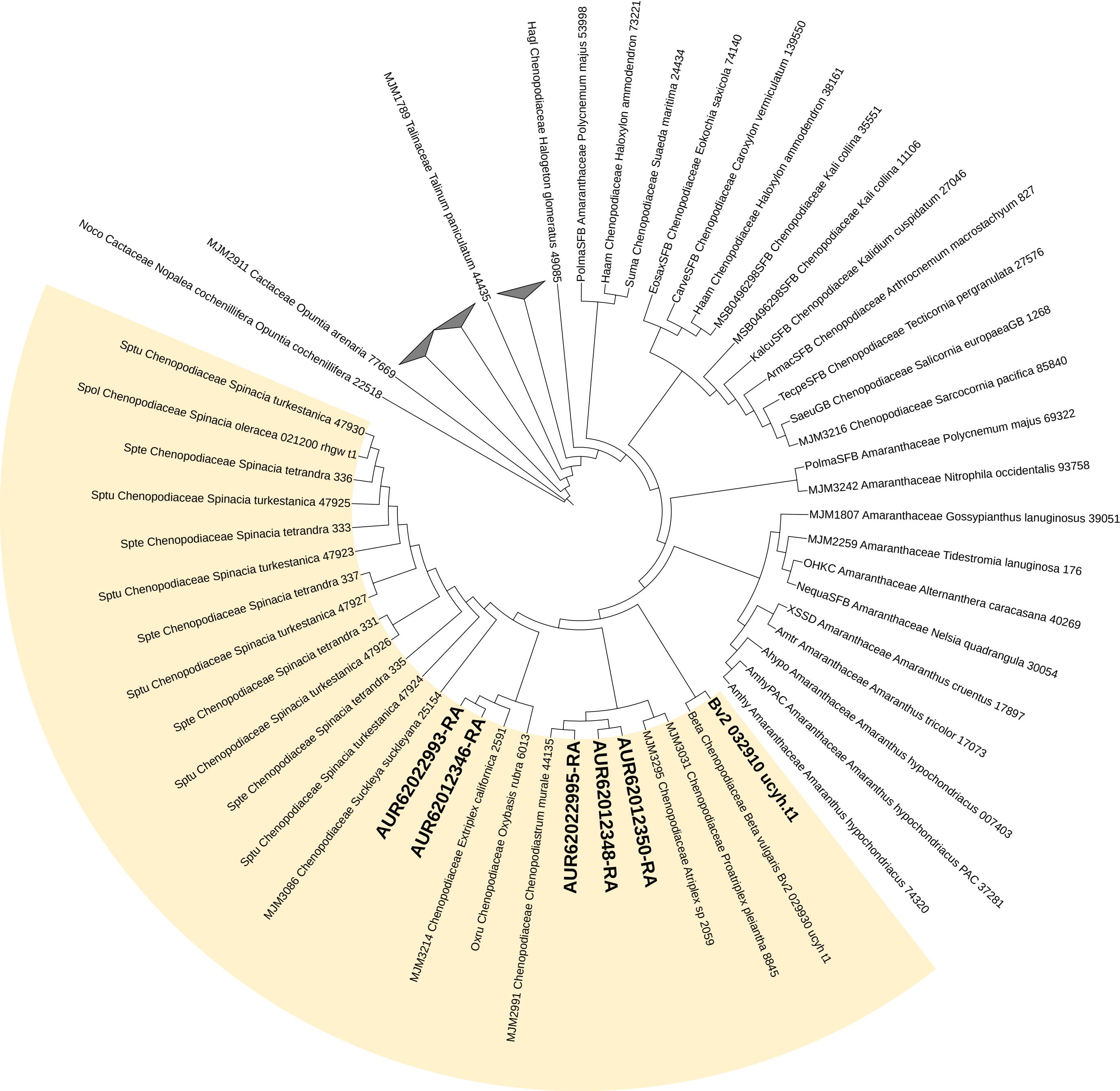

### Additional file H

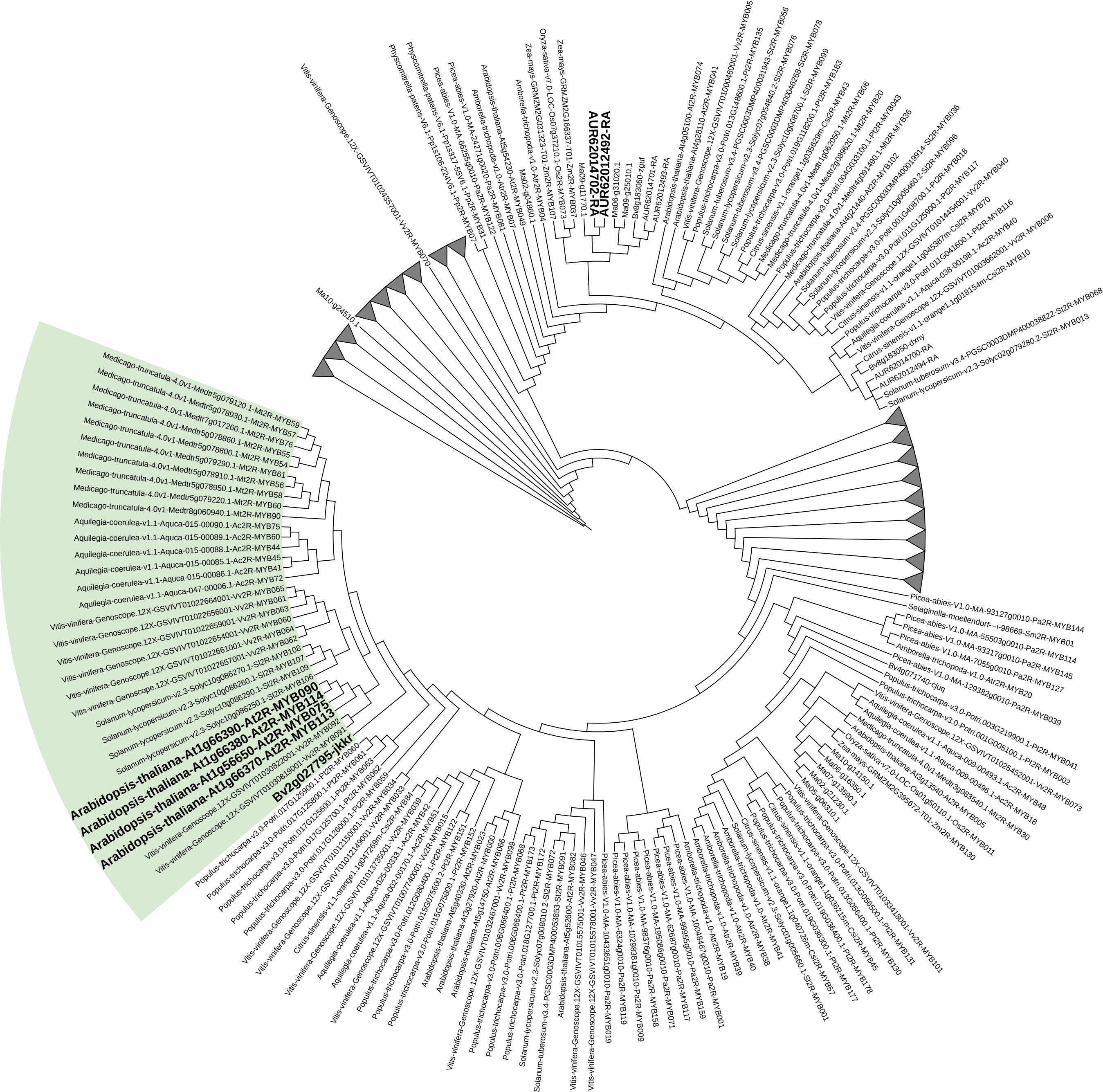
