## Additional file E for "Lack of evidence for anthocyanins contributing to pigmentation of *Chenopodium quinoa*"

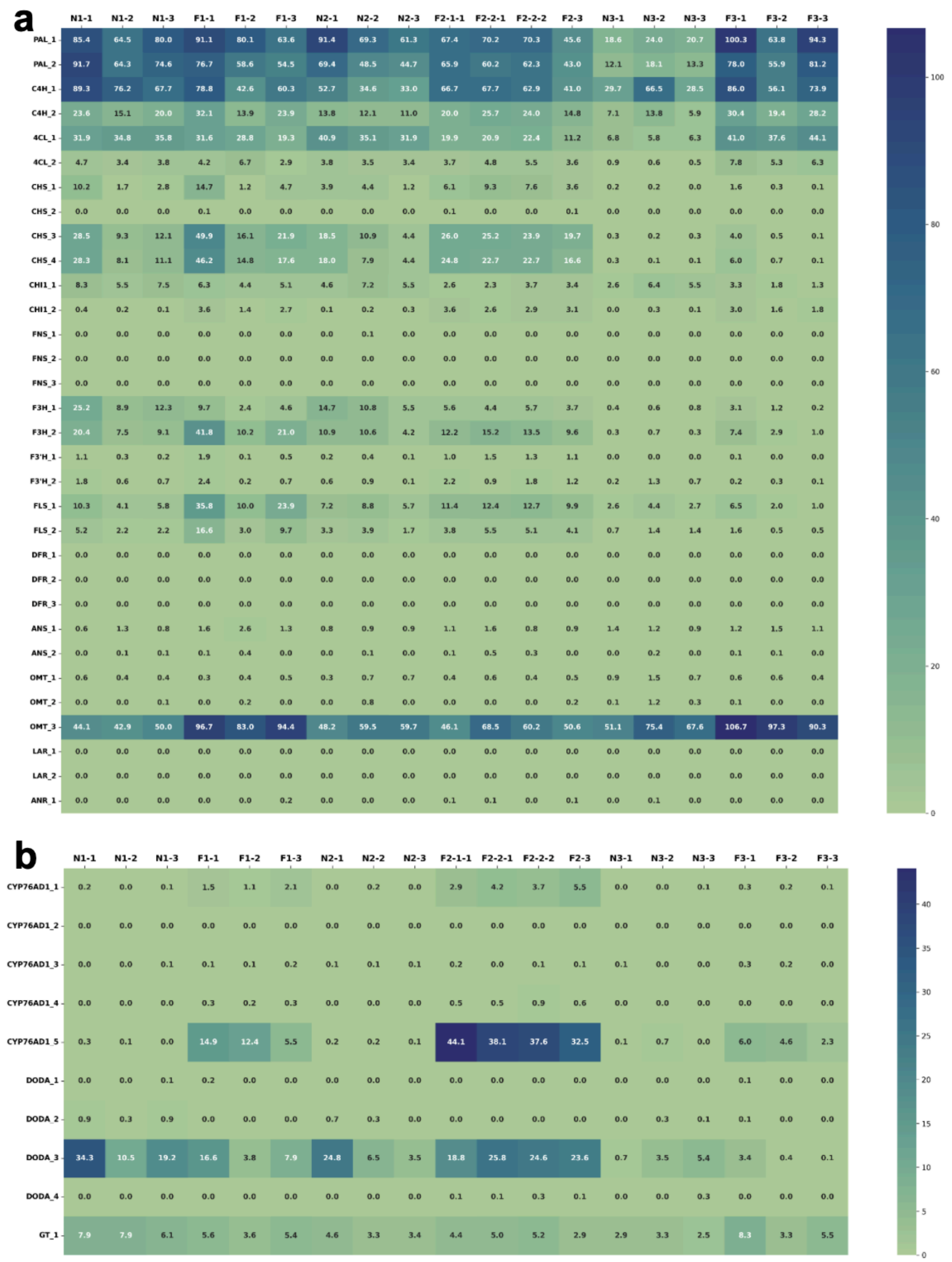


**Fig. E**: Full expression analysis for genes involved in the anthocyanin (a) and betalain (b) biosynthesis. Phenylalanine ammonia-lyase (PAL), cinnamate 4-hydroxylase (C4H), 4-coumarate-CoA ligase (4CL), chalcone synthase (CHS), chalcone isomerase (CHI), flavone synthase (FNS), flavanone 3-hydroxylase (F3H), flavonoid 3'-hydroxylase (F3'H), flavonol synthase (FLS), dihydroflavonol 4-reductase (DFR), anthocyanidin synthase (ANS), (OMT), leucoanthocyanidin reductase (LAR) and anthocyanidin reductase (ANR) are enzymes associated with the anthocyanin biosynthesis. CYP76AD1, L-DOPA 4,5-dioxygenase (DODA), UDP–glucose: cyclo-DOPA 5-O-glucosyltransferase (GT) are enzymes in the betalain biosynthesis.
